# Gene editing of pigs to control influenza A virus infections

**DOI:** 10.1101/2024.01.15.575771

**Authors:** Taeyong Kwon, Bianca L. Artiaga, Chester D. McDowell, Kristin M. Whitworth, Kevin D. Wells, Randall S. Prather, Gustavo Delhon, Mark Cigan, Stephen N. White, Jamie Retallick, Natasha N. Gaudreault, Igor Morozov, Juergen A. Richt

## Abstract

Proteolytic activation of the hemagglutinin (HA) glycoprotein by host cellular proteases is pivotal for influenza A virus (IAV) infectivity. Highly pathogenic avian influenza viruses possess the multibasic cleavage site of the HA which is cleaved by ubiquitous proteases, such as furin; in contrast, the monobasic HA motif is recognized and activated by trypsin-like proteases, such as the transmembrane serine protease 2 (TMPRSS2). Here, we aimed to determine the effects of TMPRSS2 on the replication of pandemic H1N1 and H3N2 subtype IAVs in the natural host, the pig. The use of the CRISPR/Cas 9 system led to the establishment of homozygous gene edited (GE) *TMPRSS2* knockout (KO) pigs. Delayed IAV replication was demonstrated in primary respiratory cells of KO pigs *in vitro*. IAV infection *in vivo* resulted in significant reduction of virus shedding in the upper respiratory tract, and lower virus titers and pathological lesions in the lower respiratory tract of *TMPRSS2* KO pigs as compared to WT pigs. Our findings could support the commercial use of GE pigs to minimize (i) the economic losses caused by IAV infection in pigs, and (ii) the emergence of novel IAVs with pandemic potential through genetic reassortment in the “mixing vessel”, the pig.

## Introduction

Influenza A virus (IAV) infects birds and mammals causing respiratory disease, seasonal flu, occasionally pandemics, and significant economic losses. The virus is a negative-stranded, segmented RNA virus and belongs to the family *Orthomyxoviridae*. The envelope of IAV contains two major surface glycoproteins, hemagglutinin (HA) and neuraminidase (NA), as well as in low abundance the M2 protein ^1,2^. The HA is involved in viral attachment to the cellular receptor, sialic acids, and fusion to the endosomal membrane during viral entry, whereas the NA cleaves sialic acids from host glycoproteins to facilitate the release of the newly budded virus ^3,4^. IAVs are divided into 18 different HA and 11 different NA subtypes. Avians are the natural host for all known subtypes of IAV except for H17N10 and H18N11 which are only found in bats so far ^5,6^. IAV subtypes H1N1 and H3N2 are routinely circulating in humans, resulting in seasonal flu with 290,000 to 650,000 deaths occurring every year worldwide ^7^. Human infections with avian IAV are uncommon due to different receptor specificity of human and avian IAVs and a differential distribution of these receptors in the human respiratory tract; however, infections of humans with highly pathogenic avian influenza viruses (HPAIVs) of the H5 and H7 subtypes occasionally causes fatal illness ^8^. Pigs are also an important reservoir for IAVs since they express both α2,3-linked sialic acid preferentially used by avian-like IAVs and α2,6-linked sialic acid preferentially used by mammalian-like IAVs, allowing pigs to become infected by both, mammalian- and avian-like IAVs ^9^. Therefore, pigs are considered “mixing vessels” for the generation of novel reassortant influenza viruses containing gene segments from human, swine, and/or avian IAVs ^10^. The genetic mosaic of reassortant IAVs with gene segments derived from different host origins could change the host specificity of the reassortant IAV and allow to cross species barriers. In addition, antigenic properties of the reassortant viruses might be divergent from circulating viruses, allowing these novel reassortant IAVs to evade pre-existing immunity. As a result, infection of pigs with IAVs of different origins such as avian, swine and/or human, and the subsequent genetic reassortment could have a significant impact on public health and socioeconomics, as demonstrated by the H1N1 pandemic in 2009 which originated in swine ^11^.

Based on their pathogenicity in poultry, avian IAVs can be classified into low pathogenic and highly pathogenic avian influenza viruses (LPAIV and HPAIV, respectively). The amino acid motif of the HA cleavage site is a key determinant of IAV virulence, because the cleavage site is recognized by cellular proteases which cleave the precursor protein HA0 into the HA1 and HA2 subunits, which are involved in viral attachment and entry ^12,13^. HPAI H5 and H7 subtype viruses possess a polybasic cleavage site which is efficiently cleaved by the ubiquitous subtilisin-like proteases, such as furin ^14,15^. In contrast, the monobasic cleavage motif present in the Has of LPAIVs is cleaved by trypsin-like proteases that are restricted to the respiratory and intestinal tracts of avian species ^16^. Similar to LPAIVs, mammalian-like IAVs possess a single arginine at the HA cleavage site which is cleaved by trypsin-like proteases that are present in the airways. It has been well established that the transmembrane protease serine 2 (TMPRSS2) efficiently cleaves Has of mammalian-like IAVs and plays a pivotal role in pathogenicity and infectivity of the IAVs *in vitro* and *in vivo*. Expression of TMPRSS2 allows MDCK cells to activate the monobasic HA without trypsin treatment *in vitro* ^17,18^. *In vivo* experiments showed reduced pathogenicity and virus replication of IAVs harboring the monobasic motif in *TMPRSS2* knockout (KO) mice ^19–23^.

Despite the important role of pigs in IAV evolution and ecology as the mixing vessel, gene edited (GE) pigs which are resistant or resilient to IAV infection have not been described yet. Here, we tested the hypothesis that IAV infection and transmission will be abrogated in pigs devoid of the TMPRSS2 protease. We used GE to knockout the *TMPRSS2* gene and generated homozygous *TMRPSS2* KO pigs which were tested for resistance to H1N1 and H3N2 subtype influenza virus infection.

## Results

### Generation and confirmation of gene edited TMPRSS2 KO pigs

The CRISPR/Cas-9 system was used to successfully produce *TMPRSS2* KO pigs ^24^; multiple rounds of breeding resulted in the development of a stable colony of homozygous *TMPRSS2* KO pigs which exhibit a healthy knock-out phenotype (Fig. 1A and 1B).

**Fig. 1.**
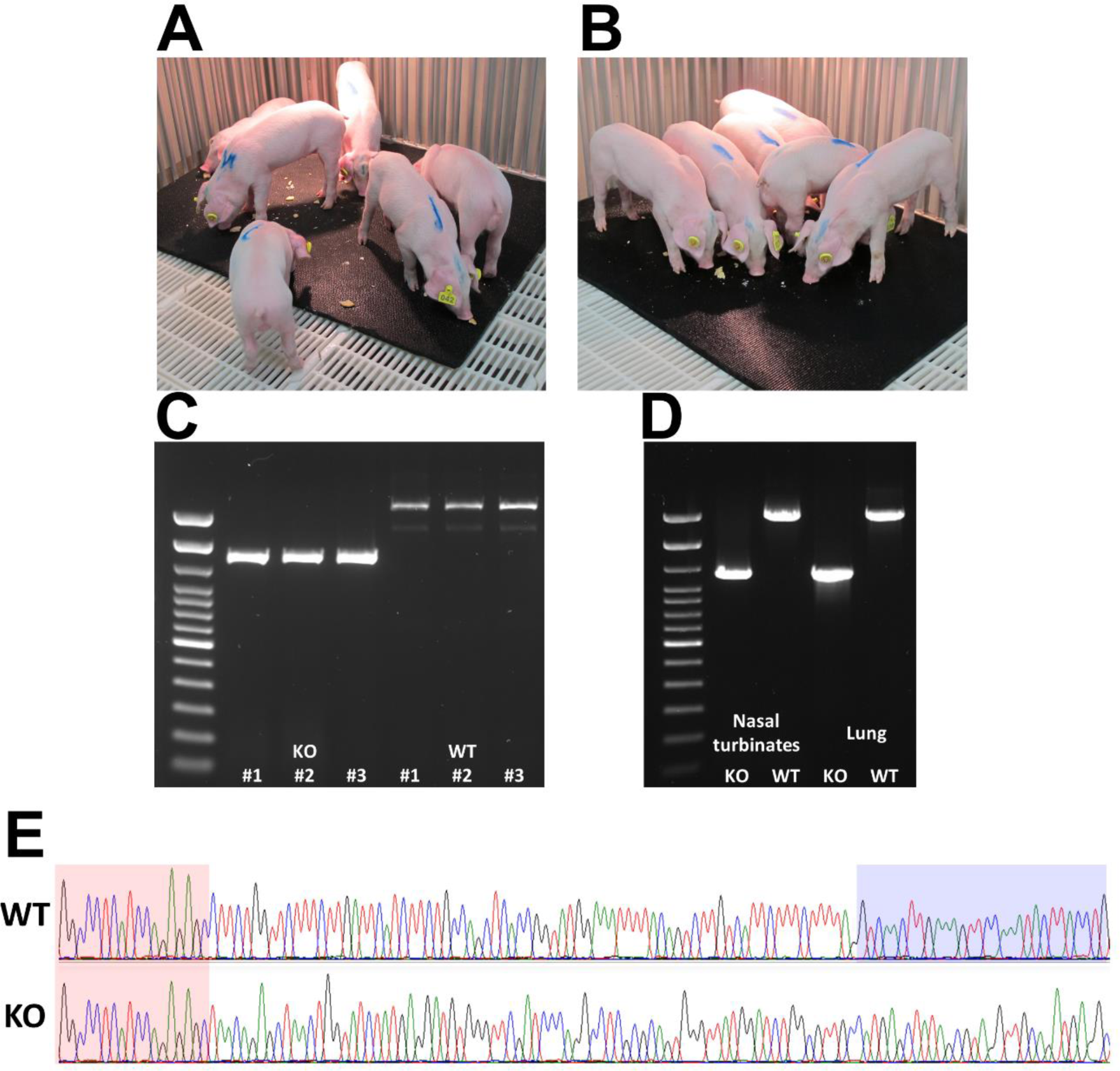
Generation and confirmation of transmembrane serine protease 2 (*TMPRSS2*) knockout (KO) pigs. Littermate *TMPRSS2* KO (A) and wild-type (WT) (B) pigs were used in this study. PCR-based genotyping confirmed *TMPRSS2* exon 2 deletion in KO pigs using tail DNA (C). *TMPRSS2* exon 2 deletion were corroborated in upper and lower respiratory tracts (D) and followed by sanger sequencing (E). Common nucleotide sequences of chromosome 13 of TMPRSS2 KO and WT pigs are shown (red shade); only DNA from WT pigs contained the exon 2-specific sequence (blue shade).

In order to confirm the knock-out of *TMPRSS2* in GE pigs, swine DNA was extracted from tails, nasal turbinates and lungs, and gene-specific primers were used to amplify the target sequence near exon 2 ^24^. In the TMPRSS2-specific PCR assay using tail samples, the PCR amplicon was 1316 bp for the KO allele, or 2181 bp for the WT allele (Fig. 1C). To further support these findings with other tissues such as respiratory tract samples, *TMPRSS2*-specific PCR was performed using homogenates of nasal turbinates and lungs, followed by Sanger sequencing. The results revealed different sizes of DNA amplicons of 1113 bp versus 1978 bp in the lungs of TMPRSS2 KO and WT pigs, respectively (Fig. 1D). In addition, the TMPRSS2 sequence of exon 2 was identified in the lungs of WT pigs, whereas the TMPRSS2 exon 2 region was absent in lungs of KO pigs (Fig. 1E). Furthermore, we confirmed the deletion of exon 2 in *TMPRSS2* mRNA transcript of KO pigs (data not shown).

### IAV infection of wild-type and TMPRSS2 KO porcine bronchial epithelial cells

To determine whether *TMPRSS2* disruption impacts susceptibility of respiratory cells to IAV infection, we isolated porcine bronchial epithelial cells (PBEC) from *TMPRSS2* KO and WT littermates for inoculation with pH1N1 (CA04 isolate) or H3N2 (TX98 isolate) *in vitro*. After 72 hours of virus infection, PBEC were methanol-fixed, and evaluated for presence of viral antigen. Using an indirect immunofluorescence assay (IFA) we could show that both KO and WT pigs were productively infected by both IAV strains. To further determine the susceptibility of PBEC to IAVs in more detail, we evaluated culture media supernatants collected at 6, 12, 24, 36, 48, and 72 hours post infection (hpi) for the presence of infectious CA04 and TX98 viruses (Fig. 2A and 2B). Infection of WT PBEC with both, pH1N1 CA04 and H3N2 TX98 viruses, resulted in the highest virus titers at 36 hours post infection; a similar titer was maintained for the remaining timepoints tested at 48 and 72 hpi. In contrast, both IAV strains presented reduced growth in *TMPRSS2* KO PBEC after 24 hpi, with H1N1 CA04 reaching WT PBEC titers at 48 hpi (Fig. 2A) and H3N2 TX98 reaching WT PBEC titers only at 72 hpi (Fig. 2B). Therefore, absence of TMPRSS2 in pig respiratory cells significantly affects IAV replication.

**Fig. 2.**
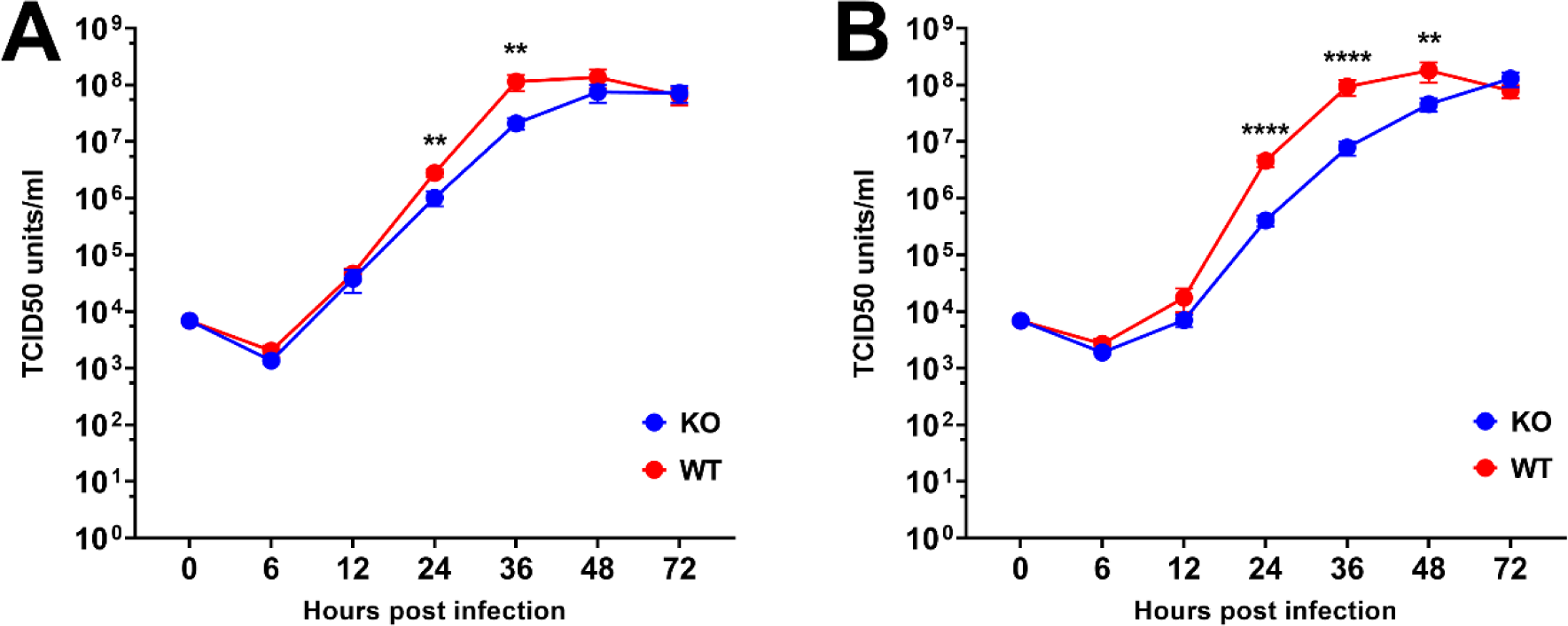
Growth curve of influenza A virus after *in vitro* infection of *TMPRSS2* KO and WT cells. Bronchial epithelial cells were collected from four *TMPRSS2* KO and three WT pigs and infected with 0.01 MOI of H1N1 CA04 (A) or H3N2 TX98 (B). Cell culture supernatant was collected at different timepoints for viral titration in MDCK cells and represented as TCID_50_/mL. Titers are represented as mean ± SEM. Statistical differences were assessed by 2-way ANOVA and Šídák’s multiple comparisons test (p-value <0.01: ** and <0.0001: ****).

### IAV infection of WT and TMPRSS2 KO pigs

In order to determine the effect of swine TMPRSS2 on IAV replication *in vivo*, littermates of homozygous TMPRSS2 KO and WT pigs were infected with either pH1N1 or H3N2 IAVs via the intra-tracheal route. Following challenge of KO pigs with pH1N1 CA04, only low levels of virus was shed via the nasal cavity at 1-day post challenge (dpc), with no virus shedding observed between 2 and 5 dpc (Fig. 3A). In contrast, virus shedding of WT pigs via the nasal cavity continued to increase from day 1 to 5 dpc (Fig. 3A), and the virus titers in nasal swab samples were statistically significantly different between the KO and WT pigs at 3, 4, and 5 dpc. When nasal turbinates were analyzed for the presence of infectious pH1N1 CA04 virus, none of KO pigs were positive at 3 and 5 dpc; in contrast, pH1N1 CA04 virus was isolated from one out of three WT pigs at 3 dpc and two out of the three WT pigs at 5 dpc (Fig. 3B). In addition, the pH1N1 CA04 virus titer in the bronchioalveolar lavage fluid (BALF) of KO pigs was lower than that in WT pigs at 5 dpc (Fig. 3C).

**Fig. 3.**
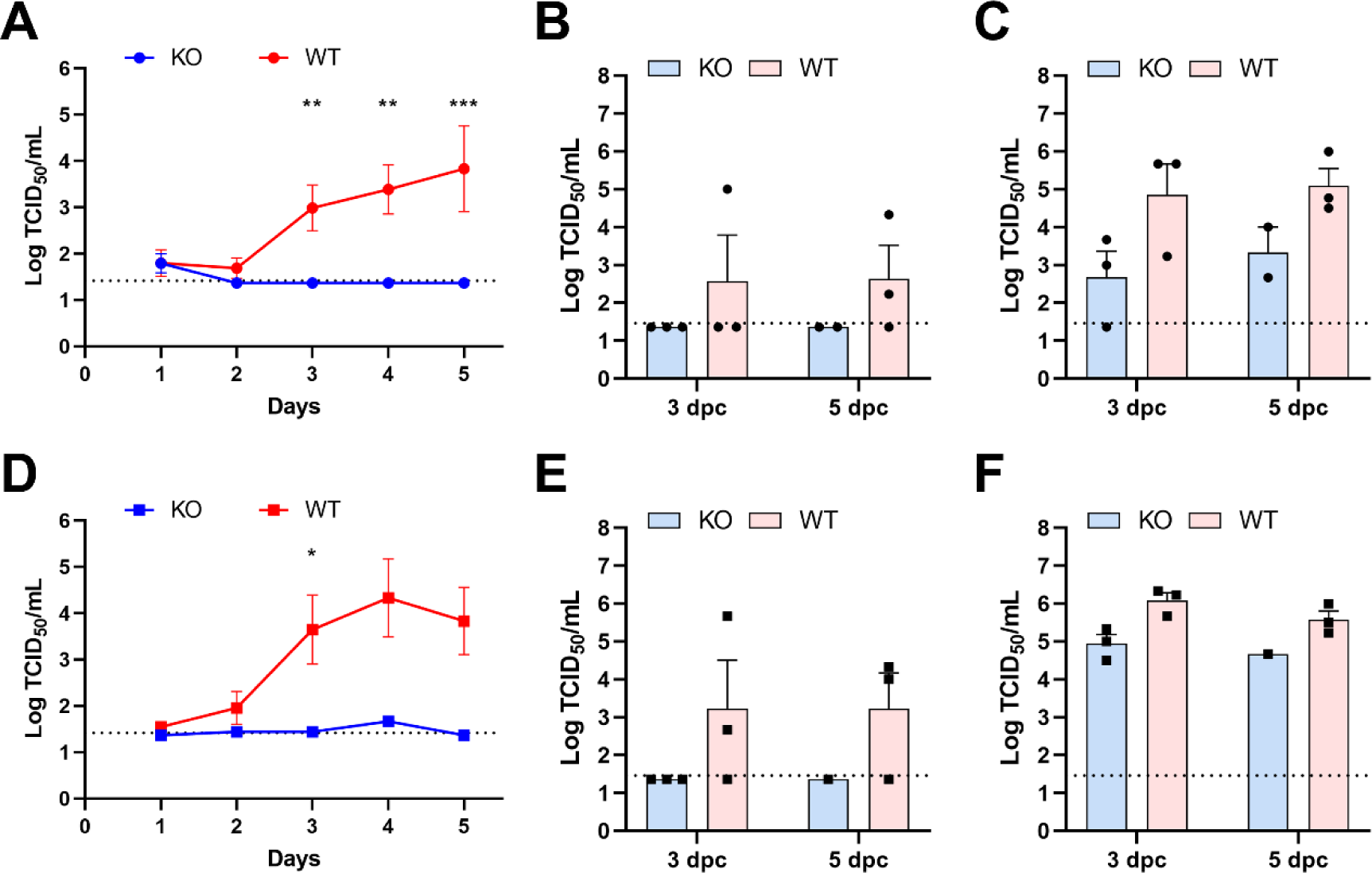
Influenza A virus shedding and replication in *TMPRSS2* KO and WT pigs. The pigs were infected with 10^6^ TCID_50_ of H1N1 CA04 (A, B, and C) or H3N2 TX98 (D, E, and F) via intra-tracheal route. Nasal swabs were collected daily, and pigs were humanly euthanized at 3- and 5-days post-challenge (dpc). Nasal swab and tissue samples were titrated on MDCK cells and represented as TCID_50_/mL. Virus titers were log-transformed for statistical analysis, and the titers are represented mean ± SEM of log-transformed values. Statistical differences were assessed by 2-way ANOVA and Šídák’s multiple comparisons test (p-value <0.05: *, <0.01: **, and <0.001: ***).

Similarly, we found minimal virus shedding from the nasal cavity of the H3N2 TX98-infected KO pigs at 2, 3 and 4 dpc (Fig. 3D). In contrast, virus shedding in WT pigs infected with the H3N2 TX98 started at 1 dpc with a peak at 4 dpc (10^4.333^ TCID_50_/mL) and still present at 5 dpc (Fig. 3D). A statistically significant difference in virus shedding via the nasal cavity between the H3N2 TX98-infected KO and WT pigs was found at 3 dpc (Fig. 3D). Importantly, no virus was present in nasal turbinates of H3N2 TX98-infected KO pigs at 3 and 5 dpc; in contrast, two out of three H3N2 TX98-infected WT pigs at 3 dpc and 5 dpc had infectious virus in nasal turbinates (Fig. 3E). In addition, the H3N2 TX98-infected KO pigs had lower virus titers than the WT pigs in the BALF at 3 dpc and at 5 dpc (Fig. 3F).

To further elucidate the viral evolution of the inoculated IAVs in the lower respiratory tract, BALF samples were subjected to next generation sequencing (NGS). The results revealed that the HA cleavage site of the pH1N1 CA04 isolate, having a IQSR motif at positions 324–327 (H1 numbering), and the H3N2 TX98 isolate, having a KQTR motif at positions 326–329 (H3 numbering), were conserved in viruses isolated from both KO and WT pigs at all time points analyzed post infection (Fig. 4). In addition, there was no amino acid (AA) substitution in the key residue “asparagine (N)” at AA position 8 of the H3N2 HA in this study: a N at this position is critical since the loss of a N-glycosylation site is known to contribute to TMPRSS2-independent activation of H3N2 subtypes in *TMPRSS2* KO mice ^25^. In addition, it was reported that a combination of an E31D substitution and the cleavage motif “GMRNVPEKQTR” (AA positions 320 to 330) resulted in efficient replication of an H3N2 virus in *TMPRSS2* KO mice ^26^. All IAV samples analyzed from TMPRSS2 and WT pigs in this study contained an N at AA position 31, and no substitutions within the cleavage motif “GMRNVPEKQTR” (<1% frequencies). The impact of N at AA position 31 on HA cleavage is unknown.

**Fig. 4.**
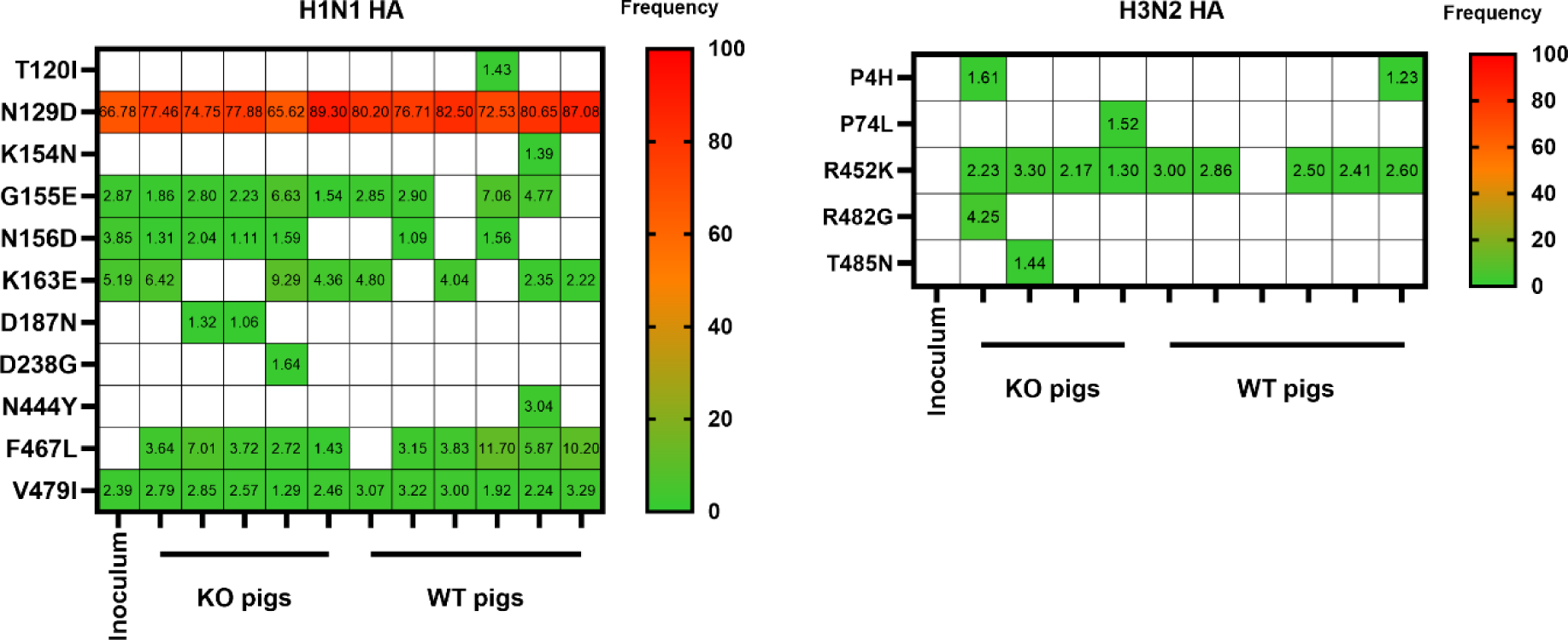
Viral evolution in bronchioalveolar fluids of KO and WT pigs. Frequencies of more than 1% of non-synonymous mutations of the HAs of H1N1 CA04 (A) and H3N2 TX98 (B) are shown.

Postmortem evaluation showed that infection of KO pigs with pH1N1 CA04 resulted in less severe macroscopic pathology of the lungs at 5 dpc when compared to WT pigs (Fig. 5A). The microscopic score of pH1N1 CA04 infected pigs generally correlated with the findings in gross pathology (Fig. 5B).

**Fig. 5.**
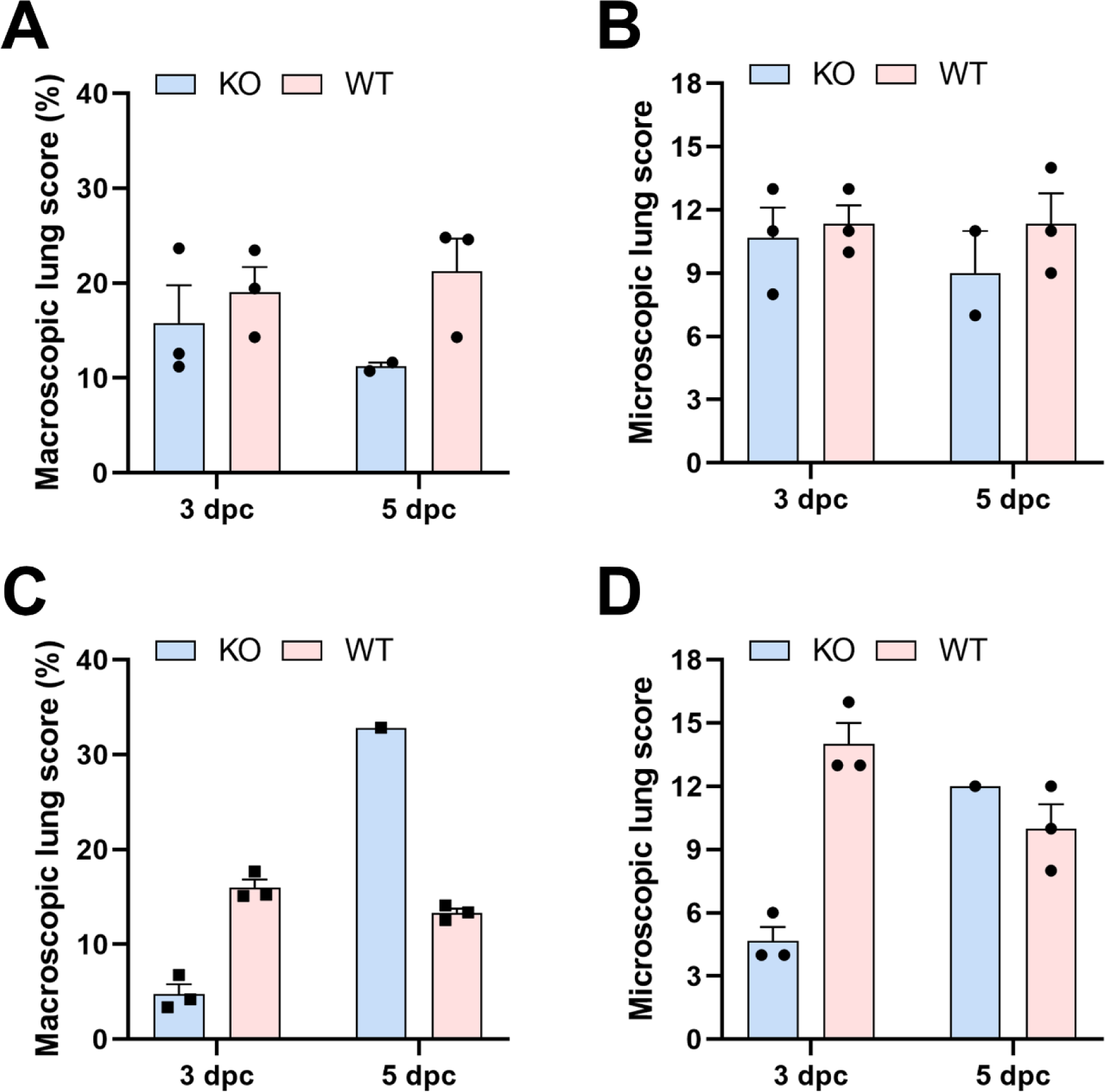
Macroscopic and microscopic lung scores in *TMPRSS2* KO and WT pigs. H1N1 CA04-(A and B) or H3N2-infected (C and D) pigs were humanly euthanized and necropsied at 3- and 5-days post-challenge (dpc). Scores are represented as mean ± SEM.

Following H3N2 TX98 infection, we found significantly less microscopic and macroscopic lung lesions in the KO pigs compared to WT pigs at 3 dpc (Fig. 5C and 5D).

## Discussion

Despite the tremendous efforts to prevent and control infectious diseases in agricultural animals using different mitigation strategies including vaccines and antiviral drugs, viruses evolve and escape vaccine-induced or naturally acquired immune responses and are able to adapt to new environments; this leads to significant challenges for vaccinology and antiviral drug development and to a significant reduction of the effectiveness and efficacy of commercial countermeasures. This challenge has offered opportunities for alternative approaches, including the generation of genetically modified livestock species that are resistant or less susceptible to infectious diseases of interest. This can be done by manipulating the host factor(s) which are critical for the virus replication cycle. The best example of gene-edited, disease-resistant pigs is the generation of *CD163* KO pigs, which have been shown to be fully resistant to porcine reproductive and respiratory syndrome virus infections ^27–29^. Other examples include the disruption of the porcine amino peptidase N (APN) or the anthrax toxin receptor 1 (ANTXR1) which resulted in resistance to infection with transmissible gastroenteritis virus or senecavirus A virus, respectively ^30,31^. In the present study, a stable colony of homozygous GE TMPRSS2 KO pigs was established. The CRISPR/Cas9 system was used to delete the start codon-containing exon 2 of the swine *TMPRSS2* gene, and this genetic change was conserved across several generations of KO animals. Moreover, molecular analysis of three different tissues (i.e., tails, nasal turbinates and lungs) confirmed that the *TMPRSS2* KO pigs were homozygous and not mosaic animals. Mosaic animals are often found in founder animals created by using the zygote injection technique for gene-editing ^32^. Furthermore, the body shape, behavior and growth of KO and WT pigs were indistinguishable. In addition, the colony of homozygous *TMPRSS2* KO pigs is maintained with no obvious anatomical alterations and reproductive issues under standard husbandry conditions (personal communication).

In the present study, the knockout of the transmembrane serine protease 2, *TMPRSS2*, and its effect on virus replication, tissue tropism and pathogenicity of two IAV subtypes, H1N1 and H3N2, was investigated both *in vitro* and *in vivo*. *In vitro* experiments showed delayed virus replication kinetics in primary bronchial epithelial cells derived from KO pigs versus WT pigs, albeit the virus titers reached a comparable peak at a late replication stage (72 hpi). Interestingly, IAV infection of KO pigs resulted in minimal virus shedding from the nasal cavity in the course of 5 days infection, whereas WT pigs shed virus continuously from 1 through 5 dpc through the nasal cavity. Furthermore, IAV infection with both IAV subtypes resulted in lower viral burden in the BALF and milder microscopic and macroscopic pathology in the KO pigs when compared to WT pigs.

Primary cell cultures are a good way to study host-virus interactions because their phenotype is closely related to that of the cells making the tissue, thereby providing a natural physico-chemical environment. Several studies have shown efficient IAV replication in swine airway explants or in swine primary respiratory epithelial cells without the treatment of exogenous trypsin, indicating the presence of endogenous proteases for cleavage and activation of the HA ^33,34^.Indeed, two swine endogenous proteases, swine *TMPRSS2* and swine airway trypsin-like protease (swAT or swTMPRSS11D), were subsequently found to be responsible for activating IAVs containing monobasic HA cleavage sites ^35^. Consistent with previous reports, we also observed efficient virus replication of pH1N1 and H3N2 IAVs in swine primary bronchial epithelial cells without treatment with exogenous trypsin. Interestingly, in primary bronchial epithelial cells derived from *TMPRSS2* KO pigs, we found delayed replication of both, pH1N1 and H3N2 IAVs at the early time points post infection; however, at late time points post infection (i.e., 72 hpi) and in terms of peak virus titers there was no difference between the *TMPRSS2* KO and the WT cells (Fig. 2A and B). Interestingly, a previous study found absence of A/Anhui/1/13 (H7N9) replication and significantly impaired replication of A/PR8/1/34 (H1N1) in tracheal, bronchial, and lung *ex vivo* explants of *TMPRSS2* KO mice, whereas replication of A/Aichi/2/68 (H3N2) was only marginally reduced in bronchial explants of *TMPRSS2* KO mice ^23^. In primary human airway cells, the knockdown of *TMPRSS2* expression by an anti-sense peptide-conjugated phosphorodiamidate morpholino oligomer that interferes with splicing of *TMPRSS2* pre-mRNA, resulted in a 2 log reduction of the virus titer of A/Anhui/1/13 (H7N9) ^36^. Here, we evaluated IAV replication in swine primary bronchial epithelial cells where not only *TMPRSS2* but also *TMPRSS11D* are expressed ^35^. Therefore, it is plausible that other proteases are able to exert their proteolytic activities for activating monobasic HAs of IAVs in the absence of T*MPRSS2*. In addition, the different host origins of the primary cells used for virus replication studies and the different IAV strains tested in these systems could account for the differences observed. The present study demonstrates altered viral replication and reduced pathogenicity of pH1N1 and H3N2 subtype IAVs in homozygous *TMPRSS2* KO pigs compared to WT littermates. Notably, infected KO pigs shed both IAVs at a low level through the nasal cavity and no virus was isolated from the nasal turbinates at 3 and 5 dpc. Our understanding on the effect of TMPRSS2 on IAV replication in the upper respiratory tract still remains unclear as evaluations of virus replication in the past mainly focused on the lower respiratory tract, i.e. lungs, which were largely performed in murine models ^19–23^. In contrast to natural hosts of IAVs (e.g., humans, swine, ferrets), there is a lack of virus shedding in experimentally IAV-infected mice, which would negate studies to determine the role of host factors in virus replication in the upper respiratory tract of mice ^37^. The present study used a natural host of IAVs, the pig, to investigate viral replication, tropism, and pathogenicity of IAVs in the upper respiratory tract of WT and TMPRSS2 KO pigs. Upon intra-tracheal challenge of pigs with swine-adapted IAVs, they generally shed virus via the nasal cavity for up to 5-7 days with a peak of virus shedding occurring at days 3 to 5 post challenge ^38–40^. This dynamic of nasal virus shedding correlates with influenza disease progression in pigs which start to recover 5 to 7 days after infection ^41,42^.. In contrast to the natural progression of disease and virus shedding in WT pigs, the GE homozygous *TMPRSS2* KO animals showed impaired replication of pH1N1 and H3N2 in nasal turbinates, which subsequently led to low virus shedding via the nasal cavity, indicating the crucial role of TMPRSS2 in IAV replication in the upper respiratory tract of swine. This indicates that TMPRSS2 plays a critical role in virus shedding and transmission of IAVs.

Another key finding of our study was the lower virus titers in BALF and less lung pathology in *TMPRSS2* KO pigs compared to WT animals, suggesting reduced virus replication, and associated pathological lesions not only in the upper but also in the lower respiratory tract. These results are consistent with previous findings in murine models where H1N1, H3N2, and H7N9 infection led to reduced lung pathology, and lower viral titers in the lungs of *TMPRSS2* KO compared to WT mice ^19,22,23^. In addition, reassorted IAVs harboring an avian H2 or H10 HA gene segment in the A/Puerto Rico/8/34 (PR8 H1N1 virus) backbone were not replicating efficiently in the lower respiratory tract of *TMPRSS2* KO mice ^20,21^. While mice are a well-established animal model for influenza virus studies, they are not natural hosts of IAVs and it requires serial passages for virus adaptation^37,42^. In contrast, pigs are a natural host for IAV infections, thus they might better represent a human-like host-pathogen interactions during the course of IAV infection. In this respect, our study confirmed the effect of TMPRSS2 on the replication and pathogenicity of IAVs in the upper and lower respiratory tracts of a natural host species. Both pH1N1 and H3N2 subtypes were only shed in low amount for the nasal cavity although they were able to replicate in the lungs of *TMPRSS2* KO pigs, however to lower virus titers and less pathological lesions when compared to WT control animals. This indicates that pH1N1 and H3N2 subtype IAVs may utilize TMPRSS2-independent activation of the HA in the lower respiratory tract of pigs. Cells in the lower respiratory tract not only express a variety transmembrane serine proteases, such as TMPRSS2, TMPRSS4 and TMPRSS11D, but they also secrete soluble proteases into airways, such as tryptase Clara, all of which have been shown to be involved in proteolytic cleavage of monobasic HAs of IAVs ^17,18,43^. Importantly, we found that the monobasic HA cleavage sites as well as other key residues determining efficient HA cleavage in IAVs replicating in *TMPRSS2* KO pigs remained unchanged in IAVs replicating for several days in both, the *TMPRSS2* KO and WT pigs, suggesting that there no evidence of emerging viral variants exhibiting potentially different geno-and phenotypes for HA activation.

Another example of gene editing of animals for IAV resistance is the *ANP32A* modification in chickens with two amino acid substitutions, N129I and D130N ^44^. Human ANP32A and ANP32B are crucial for influenza virus replication through the formation of functional influenza replicases ^45,46^. In chickens, only ANP32A is responsible for the activation of the influenza polymerase complex; this is attributed to amino acid differences at positions of 129 and 130 in chicken ANP32A and ANP32B(129N and 130D in chicken ANP32A, 129I and 130N in chicken ANP32B) ^47,48^. It was recently shown that experimental infection of GE ANP32A chickens with a low dose of an H9N2 avian influenza virus resulted in one out of 10 chickens shedding virus via the oropharyngeal route; and the virus was not transmitted to 10 in-contact GE APN32A chickens ^44^. Resilience of the GE APN32A chickens to high-dose influenza virus infection was supported by: (1) inefficient transmission from principal-infected WT chickens to one out of four in-contact GE chickens, (2) a low level of virus shedding in five out of 10 principal-infected GE chickens, and (3) no transmission from principal-infected GE chickens to in-contact GE chickens ^44^. Importantly, critical AA substitutions in two polymerase genes, such as PB2-M631L and PA-E349K, were observed after infection; these AA changes might be responsible for the emergence of escape mutants in the GE APN32A chickens^44^.

In summary, our study elucidates the effect of the TMPRSS2 protease on IAV replication in a natural host species, the pig. To this end, we generated homozygous *TMPRSS2* KO pigs by using the CRISPR/Cas9 technology and established a stable and healthy *TMPRSS2* KO colony. *In vitro* experiments showed delayed virus replication in primary respiratory KO cells compared to WT controls. In *in vivo* experiments, a significant reduction of virus shedding via the nasal cavity and lower virus titers and pathological lesions in the lower respiratory tract of *TMPRSS2* KO pigs as compared to WT pigs was demonstrated. The effect of the *TMPRSS2* deletion on IAV replication and nasal shedding in pigs support the notion of a commercial use of such GE swine in order to minimize (i) the economic losses caused by swine influenza virus infection for the pork industry, and (ii) the risk of the emergence of novel influenza viruses with pandemic potential through genetic reassortment in the “mixing vessel”, the pig.

## Methods

### Cells and viruses

Madin-Darby canine kidney (MDCK) cells were maintained in Dulbecco’s Modified Eagle Medium (DMEM; Corning, Manassas, VA, USA) supplemented with 5% fetal bovine serum (FBS; R&D systems, Flower Branch, GA, USA) and 1% antibiotic-antimycotic (Gibco, Grand Island, NY, USA). The viruses used in this study were A/Swine/Texas/4199-2/98 (H3N2) (TX98 H3N2) and A/California/04/2009 (H1N1) (CA04 pH1N1) ^49,50^. The viruses were propagated in MDCK cells in the presence of L-(tosylamido-2-phenyl) ethyl chloromethyl ketone (TPCK)-treated trypsin (Worthington Biochemical Corporation, Lakewood, NJ).

### Generation and confirmation of KO pigs

*TMPRSS2* KO pigs (RRID NSRRC:0060) were generated using the CRISPR/Cas-9 technology as previously described ^24^. Briefly, immature oocytes were collected from pre-pubertal gilts and cultured with semen for *in vitro* fertilization. At 14 hours post-fertilization, a pair of guide RNAs targeting the exon 2 of the swine *TMPRSS2* gene and polyadenylated Cas9 endonuclease were directly injected into the cytoplasm of zygotes. The modified embryos were then surgically transferred into surrogate gilts for gestation. Subsequently, the resulting founder males and females were bred to create offspring that inherited homozygotic KO (TMPRSS2 −/−), heterozygotic KO (TMPRSS2 +/−) or wild type (WT).

To screen and confirm the knock-out of the *TMPRSS2* gene, tails, nasal turbinate, and lungs of each piglet were subject to polymerase chain reaction (PCR) using gene-specific primers. Briefly, swine DNA of tails was extracted using an automatic commercial magnetic bead extraction kit (GeneReach USA, Lexington, MA). PCR was performed with medium range primers ^24^ using Q5 High-Fidelity DNA polymerase (New England Biolabs, Ipswich, MA) according to the manufacturer’s protocol. To corroborate the complete gene knockout of the exon 2 in respiratory tracts, swine DNA was extracted from nasal turbinates and lungs using DNeasy Blood & Tissue kit (Qiagen, Germantown, MD, USA) and subjected to PCR with primers (Forward primer: CCTGGATCTGAGGGAGCGA and reverse primer: AGGGCAGCGCGATTAGAAAG) using a Platinum™ Taq DNA High Fidelity polymerase (Invitrogen, Carlsbad, CA) or Platinum™ SuperFi II Green PCR Master Mix (Invitrogen, Carlsbad, CA). The amplicons of lungs of both KO and WT pigs were submitted for Sanger sequencing.

### Porcine bronchial epithelial cell (PBEC) isolation and culture

The trachea and lungs from four *TMPRSS2* KO and three WT littermates were collected after humane euthanasia. The trachea was filled with isolation medium (DMEM:F12 [Gibco, Grand Island, NY, USA] and antibiotic-antimycotic [Gibco, Grand Island, NY, USA]) and transported on ice for epithelial cell isolation starting less than 6 hours after necropsy. The bronchi tree was dissected from lung tissue and rinsed with isolation medium supplemented with 5 mM DTT (Sigma-Aldrich, St. Louis, MO) to remove mucus and debris. Bronchi were cut into 5 mm strips and incubated in isolation medium with 1.4 mg/mL Pronase (Sigma-Aldrich, St. Louis, MO) and 100 μg/ml DNase I (Sigma-Aldrich, St. Louis, MO) for 24 hours at 4°C with continuous gentle agitation. After incubation, FBS was added to inhibit protease activity. Epithelial cells were collected after vigorous agitation and lightly scraping of the tissue strips. After rinsing tissue strips with isolation medium, samples were filtered through a 70 to 100 μm cell strainer, centrifuged (200g, 10 min, 4°C), and resuspended in isolation medium with 10% FBS. Fibroblasts were removed by incubating the samples in a non-treated tissue culture dish for 2 hours at 37°C and 5% CO_2_. Non-adherent cells were centrifuged again and resuspended in 2 mL of bronchial epithelial cell growth medium (BEGM) (Lonza cat# CC-3170) supplemented with Primocin (InvivoGen, San Diego, CA). Live cells were seeded at 1×10^5^ cells/cm^2^ into collagen coated culture flasks in BEGM with Primocin and 5% FBS and incubated at 37°C and 5% CO_2_ to observe cell viability and bacterial contamination. Medium was changed every 48 hours and FBS was gradually removed from the medium composition to reduce fibroblast growth. Cell monolayer was subcultured at 80–90% confluence.

### PBEC infection with influenza A virus

After two passages in T25 flasks, PBEC were seeded into 24-well plates at 4×10^5^ cells/well. After 18 hours, BEGM was removed and the confluent monolayer was washed once with PBEC infection medium (DMEM:F12 supplemented with Primocin and 0.3% bovine albumin serum and 1% MEM vitamin). CA04 pH1N1 or TX98 H3N2 were diluted in infection medium to inoculate PBEC at 0.01 MOI in three replicates per pig. Supernatant was collected at 6, 12, 24, 36, 48, and 72 hours post infection and titrated to define the growth curve of both IAV strains in *TMPRSS2* KO and WT PBEC.

### Animal study

Twenty-one 4-week-old influenza H1 and H3 subtypes-seronegative piglets (9 KO pigs and 12 WT pigs) were used for this study. KO and WT pigs were littermates from several sows. The PCR-based genotyping of TMPRSS2 confirmed homozygous KO and WT genotypes of the pigs. The pigs were divided into two groups: five KO pigs and six WT pigs for pH1N1 CA04 infection, and four KO pigs and six WT pigs for H3N2 TX98 infection. The pigs were challenged with 10^6^ TCID_50_ of either CA04 pH1N1 or TX98 H3N2 via intra-tracheal route. Nasal swabs were collected daily in 2 mL of medium, filtered through 0.45 um syringe filter (TPP, Trasadingen, Switzerland), and stored at −80°C until further analysis. The pigs were humanely euthanized at 3 or 5 days post-challenge (dpc) for pathological evaluation and tissue sample collection. The bronchioalveolar lavage fluid was collected from the left side of the lung with 50 mL of DMEM.

### Pathology and histopathology

At necropsy, each lobe of the lung was scored to assess macroscopic lung lesions ^49^. The nasal turbinates, trachea, and right cardiac lobe were fixed in 10% buffered formalin for further histopathological evaluation. The tissues were then embedded in paraffin and stained with hematoxylin and eosin.

Microscopic lesions were blindly evaluated by a veterinary pathologist with scoring of 0 to 3 for each of six criteria: (i) epithelial necrosis, attenuation and disruption, (ii) airway exudate-necrosis and inflammation, (iii) percentage of airways with inflammation, (iv) peribronchiolar and perivascular lymphocytic inflammation, (v) alveolar exudate, and (vi) alveolar septal inflammation ^51^. The total sum of the scores was calculated for each pig (0 to 18).

### Virus titration

MDCK cells were seeded in a 96-wells plate a day before the assay to reach 90 to 100% confluence at the day of assay. Ten-fold dilutions of nasal swab sample and BALF were prepared in virus maintenance medium (DMEM supplemented with 0.3% bovine albumin serum, 1% MEM vitamin, 1% antibiotic-antimycotic, and 1 μg/mL TPCK-treated trypsin). The 10% tissue homogenate was prepared and filtered through a 0.45% syringe filter. Ten-fold dilutions of 10% tissue homogenates were also prepared in virus maintenance medium. MDCK cells were washed with phosphate buffered saline once, and virus inoculum was transferred onto the cells. For tissue homogenates, the inoculum in the first three rows were changed with fresh virus maintenance medium after 2 hours absorption at 37°C. At 48 hours post-inoculation, the cells were fixed with 100% cold methanol and incubated in −20 °C for 10 minutes. The cells were incubated with mouse monoclonal antibodies against IAV NP (HB65), followed by Alex Fluor 488 antibody (Invitrogen, Carlsbad, CA) for visualization. The virus titer was calculated using Reed-Muench methods.

### Next-generation sequencing

The whole genome sequence of CA04 pH1N1 and TX98 H3N2 was determined using an Illumina NextSeq platform. Briefly, viral RNA was extracted from challenge materials and BALF samples using a QIAamp Viral RNA mini kit (Qiagen, Germantown, MD, USA) and subjected to multi-segment reverse transcription polymerase chain reaction (RT-PCR) ^52,53^. Briefly, extracted RNA was mixed with either Uni or Opti primer sets in a Platinum SuperScript III One Step RT-PCR kits (Invitrogen, Carlsbad, CA) for amplification. The amplicons were visualized by gel electrophoresis, followed by library preparation using the Illumina DNA Prep kit (Illumina, San Diego, CA) according to manufacturer’s protocols. The libraries were then sequenced with the Illumina NextSeq platform using 150bp paired-end reads with a mid-output kit. Reads were then demultiplexed and parsed into individual FASTQ files that were imported into CLC Genomics Workbench version 22.0.1 (Qiagen, Germantown, MD, USA) for analysis. Reads were trimmed to remove primer sequences and filtered to remove low quality and short reads. The reads were mapped to the reference genome obtained from NCBI (CA04: MN371610–MN371617 and Tx98: CY095672– CY095679). Following read mapping, all samples were run through the low frequency variant caller module within CLC Genomic Workbench with a frequency cutoff greater than 1%. Heatmaps for each segment of the genome were generated for nonsynonymous variants greater than 1% using R Studio (2022.12.0+353).

## Acknowledgments

We gratefully thank Dashzeveg Bold, Cassidy Keating, Yonghai Li, Antoinette Lona, Daniel Madden, Jayme A. Souza-Neto, and Baolin Wang for assistance with the animal studies, Codie Durfee for technical assistance, Melissa Rohrer for managing the project, and the UNL animal facility personnel for assistance in maintaining the *TMPRSS2* KO pig colony. Funding for this study was provided through grants from Genus PLC, the National Bio and Agro-Defense Facility (NBAF) Transition Fund from the State of Kansas, the AMP and MCB Core of the Center on Emerging and Zoonotic Infectious Diseases (CEZID) of the National Institutes of General Medical Sciences under award number P20GM130448, the NIAID Centers of Excellence for Influenza Research and Surveillance (CEIRS) under contract number HHSN 272201400006C, and the NIAID supported Center of Excellence for Influenza Research and Response (CEIRR) under contract number 75N93021C00016. Funding for the National Swine Resource and Research Center is from the National Institute of Allergy and Infectious Disease, the National Heart, Lung and Blood Institute, and the Office of Research Infrastructure Programs, Office of the Director (U42OD011140).

## Author contributions

J.A.R designed research; T.K., B.L.A., C.D.M., K.M.W., K.D.W., R.S.P., G.D., J.R., N.N.G., and I.M. performed research and analyzed data; M.C. and S.N.W. performed project administration and data analysis; T.K. and J.A.R. wrote the paper. All authors have read and agreed to the published version of the manuscript.

## References

1 Harris, A. et al. Influenza virus pleiomorphy characterized by cryoelectron tomography. Proc Natl Acad Sci U S A 103, 19123–19127 (2006). 10.1073/pnas.0607614103

2 Zebedee, S. L. & Lamb, R. A. Influenza A virus M2 protein: monoclonal antibody restriction of virus growth and detection of M2 in virions. J Virol 62, 2762–2772 (1988). 10.1128/JVI.62.8.2762-2772.1988

3 Palese, P., Tobita, K., Ueda, M. & Compans, R. W. Characterization of temperature sensitive influenza virus mutants defective in neuraminidase. Virology 61, 397–410 (1974). 10.1016/0042-6822(74)90276-1

4 Weis, W. et al. Structure of the influenza virus haemagglutinin complexed with its receptor, sialic acid. Nature 333, 426–431 (1988). 10.1038/333426a0

5 Tong, S. et al. A distinct lineage of influenza A virus from bats. Proc Natl Acad Sci U S A 109, 4269–4274 (2012). 10.1073/pnas.1116200109

6 Tong, S. et al. New world bats harbor diverse influenza A viruses. PLoS Pathog 9, e1003657 (2013). 10.1371/journal.ppat.1003657

7 Iuliano, A. D. et al. Estimates of global seasonal influenza-associated respiratory mortality: a modelling study. Lancet 391, 1285–1300 (2018). 10.1016/S0140-6736(17)33293-2

8 Shinya, K. et al. Avian flu: influenza virus receptors in the human airway. Nature 440, 435–436 (2006). 10.1038/440435a

9 Ito, T. et al. Molecular basis for the generation in pigs of influenza A viruses with pandemic potential. J Virol 72, 7367–7373 (1998). 10.1128/JVI.72.9.7367-7373.1998

10 Scholtissek, C. Molecular evolution of influenza viruses. Virus Genes 11, 209–215 (1995). 10.1007/BF01728660

11 Mena, I. et al. Origins of the 2009 H1N1 influenza pandemic in swine in Mexico. Elife 5 (2016). 10.7554/eLife.16777

12 Garten, W. & Klenk, H.-D. Understanding influenza virus pathogenicity. Trends in Microbiology 7, 99–100 (1999). 10.1016/s0966-842x(99)01460-2

13 Horimoto, T. & Kawaoka, Y. Reverse genetics provides direct evidence for a correlation of hemagglutinin cleavability and virulence of an avian influenza A virus. J Virol 68, 3120–3128 (1994). 10.1128/JVI.68.5.3120-3128.1994

14 Stieneke-Grober, A. et al. Influenza virus hemagglutinin with multibasic cleavage site is activated by furin, a subtilisin-like endoprotease. EMBO J 11, 2407–2414 (1992). 10.1002/j.1460-2075.1992.tb05305.x

15 Vey, M. et al. Hemagglutinin activation of pathogenic avian influenza viruses of serotype H7 requires the protease recognition motif R-X-K/R-R. Virology 188, 408–413 (1992). 10.1016/0042-6822(92)90775-k

16 Bottcher-Friebertshauser, E., Garten, W., Matrosovich, M. & Klenk, H. D. The hemagglutinin: a determinant of pathogenicity. Curr Top Microbiol Immunol 385, 3–34 (2014). 10.1007/82_2014_384

17 Bottcher, E. et al. Proteolytic activation of influenza viruses by serine proteases TMPRSS2 and HAT from human airway epithelium. J Virol 80, 9896–9898 (2006). 10.1128/JVI.01118-06

18 Chaipan, C. et al. Proteolytic activation of the 1918 influenza virus hemagglutinin. J Virol 83, 3200–3211 (2009). 10.1128/JVI.02205-08

19 Hatesuer, B. et al. Tmprss2 is essential for influenza H1N1 virus pathogenesis in mice. PLoS Pathog 9, e1003774 (2013). 10.1371/journal.ppat.1003774

20 Lambertz, R. L. O. et al. H2 influenza A virus is not pathogenic in Tmprss2 knock-out mice. Virol J 17, 56 (2020). 10.1186/s12985-020-01323-z

21 Lambertz, R. L. O. et al. Tmprss2 knock-out mice are resistant to H10 influenza A virus pathogenesis. J Gen Virol 100, 1073–1078 (2019). 10.1099/jgv.0.001274

22 Sakai, K. et al. The host protease TMPRSS2 plays a major role in in vivo replication of emerging H7N9 and seasonal influenza viruses. J Virol 88, 5608–5616 (2014). 10.1128/JVI.03677-13

23 Tarnow, C. et al. TMPRSS2 is a host factor that is essential for pneumotropism and pathogenicity of H7N9 influenza A virus in mice. J Virol 88, 4744–4751 (2014). 10.1128/JVI.03799-13

24 Whitworth, K. M. et al. Zygote injection of CRISPR/Cas9 RNA successfully modifies the target gene without delaying blastocyst development or altering the sex ratio in pigs. Transgenic Res 26, 97–107 (2017). 10.1007/s11248-016-9989-6

25 Sakai, K. et al. A mutant H3N2 influenza virus uses an alternative activation mechanism in TMPRSS2 knockout mice by loss of an oligosaccharide in the hemagglutinin stalk region. J Virol 89, 5154–5158 (2015). 10.1128/JVI.00124-15

26 Lambertz, R. L. O. et al. Exchange of amino acids in the H1-haemagglutinin to H3 residues is required for efficient influenza A virus replication and pathology in Tmprss2 knock-out mice. J Gen Virol 99, 1187–1198 (2018). 10.1099/jgv.0.001128

27 Burkard, C. et al. Pigs Lacking the Scavenger Receptor Cysteine-Rich Domain 5 of CD163 Are Resistant to Porcine Reproductive and Respiratory Syndrome Virus 1 Infection. J Virol 92 (2018). 10.1128/JVI.00415-18

28 Whitworth, K. M. et al. Gene-edited pigs are protected from porcine reproductive and respiratory syndrome virus. Nat Biotechnol 34, 20–22 (2016). 10.1038/nbt.3434

29 Yang, H. et al. CD163 knockout pigs are fully resistant to highly pathogenic porcine reproductive and respiratory syndrome virus. Antiviral Res 151, 63–70 (2018). 10.1016/j.antiviral.2018.01.004

30 Chen, P. R. et al. Disruption of anthrax toxin receptor 1 in pigs leads to a rare disease phenotype and protection from senecavirus A infection. Sci Rep 12, 5009 (2022). 10.1038/s41598-022-09123-x

31 Whitworth, K. M. et al. Resistance to coronavirus infection in amino peptidase N-deficient pigs. Transgenic Res 28, 21–32 (2019). 10.1007/s11248-018-0100-3

32 Watanabe, M. & Nagashima, H. Genome Editing of Pig. Methods Mol Biol 1630, 121–139 (2017). 10.1007/978-1-4939-7128-2_11

33 Bateman, A. C., Karasin, A. I. & Olsen, C. W. Differentiated swine airway epithelial cell cultures for the investigation of influenza A virus infection and replication. Influenza Other Respir Viruses 7, 139–150 (2013). 10.1111/j.1750-2659.2012.00371.x

34 Van Poucke, S. G., Nicholls, J. M., Nauwynck, H. J. & Van Reeth, K. Replication of avian, human and swine influenza viruses in porcine respiratory explants and association with sialic acid distribution. Virol J 7, 38 (2010). 10.1186/1743-422X-7-38

35 Peitsch, C., Klenk, H. D., Garten, W. & Bottcher-Friebertshauser, E. Activation of influenza A viruses by host proteases from swine airway epithelium. J Virol 88, 282–291 (2014). 10.1128/JVI.01635-13

36 Limburg, H. et al. TMPRSS2 Is the Major Activating Protease of Influenza A Virus in Primary Human Airway Cells and Influenza B Virus in Human Type II Pneumocytes. J Virol 93 (2019). 10.1128/JVI.00649-19

37 Bouvier, N. M. & Lowen, A. C. Animal Models for Influenza Virus Pathogenesis and Transmission. Viruses 2, 1530–1563 (2010). 10.3390/v20801530

38 Ma, W. et al. Identification and characterization of a highly virulent triple reassortant H1N1 swine influenza virus in the United States. Virus Genes 40, 28–36 (2010). 10.1007/s11262-009-0413-7

39 Vincent, A. L. et al. Evaluation of hemagglutinin subtype 1 swine influenza viruses from the United States. Vet Microbiol 118, 212–222 (2006). 10.1016/j.vetmic.2006.07.017

40 Vincent, A. L. et al. Characterization of a newly emerged genetic cluster of H1N1 and H1N2 swine influenza virus in the United States. Virus Genes 39, 176–185 (2009). 10.1007/s11262-009-0386-6

41 Janke, B. H. Clinicopathological features of Swine influenza. Curr Top Microbiol Immunol 370, 69–83 (2013). 10.1007/82_2013_308

42 Rajao, D. S. & Vincent, A. L. Swine as a model for influenza A virus infection and immunity. ILAR J 56, 44–52 (2015). 10.1093/ilar/ilv002

43 Kido, H. et al. Isolation and characterization of a novel trypsin-like protease found in rat bronchiolar epithelial Clara cells. A possible activator of the viral fusion glycoprotein. Journal of Biological Chemistry 267, 13573–13579 (1992). 10.1016/s0021-9258(18)42250-8

44 Idoko-Akoh, A. et al. Creating resistance to avian influenza infection through genome editing of the ANP32 gene family. Nat Commun 14, 6136 (2023). 10.1038/s41467-023-41476-3

45 Carrique, L. et al. Host ANP32A mediates the assembly of the influenza virus replicase.Nature 587, 638–643 (2020). 10.1038/s41586-020-2927-z

46 Staller, E. et al. ANP32 Proteins Are Essential for Influenza Virus Replication in Human Cells. J Virol 93 (2019). 10.1128/JVI.00217-19

47 Long, J. S. et al. Species specific differences in use of ANP32 proteins by influenza A virus. Elife 8 (2019). 10.7554/eLife.45066

48 Zhang, H. et al. Fundamental Contribution and Host Range Determination of ANP32A and ANP32B in Influenza A Virus Polymerase Activity. J Virol 93 (2019). 10.1128/JVI.00174-19

49 Artiaga, B. L. et al. Evaluating alpha-galactosylceramide as an adjuvant for live attenuated influenza vaccines in pigs. Anim Dis 2, 19 (2022). 10.1186/s44149-022-00051-x

50 Richt, J. A. et al. Pathogenic and antigenic properties of phylogenetically distinct reassortant H3N2 swine influenza viruses cocirculating in the United States. J Clin Microbiol 41, 3198–3205 (2003). 10.1128/JCM.41.7.3198-3205.2003

51 Khurana, S. et al. Vaccine-induced anti-HA2 antibodies promote virus fusion and enhance influenza virus respiratory disease. Sci Transl Med 5, 200ra114 (2013). 10.1126/scitranslmed.3006366

52 Lee, D. H. Complete Genome Sequencing of Influenza A Viruses Using Next-Generation Sequencing. Methods Mol Biol 2123, 69–79 (2020). 10.1007/978-1-0716-0346-8_6

53 Zhou, B. & Wentworth, D. E. Influenza A virus molecular virology techniques. Methods Mol Biol 865, 175–192 (2012). 10.1007/978-1-61779-621-0_11

